# Thiocillin and Micrococcin Exploit the Ferrioxamine Receptor of *Pseudomonas aeruginosa* for Uptake

**DOI:** 10.1101/2020.04.23.057471

**Authors:** Derek C. K. Chan, Lori L. Burrows

## Abstract

Thiopeptides are a class of Gram-positive antibiotics that inhibit protein synthesis. They have been underutilized as therapeutics due to solubility issues, poor bioavailability, and lack of activity against Gram-negative pathogens. We discovered recently that a member of this family, thiostrepton, has activity against *Pseudomonas aeruginosa and Acinetobacter baumannii* under iron-limiting conditions. Thiostrepton uses pyoverdine siderophore receptors to cross the outer membrane, and combining thiostrepton with an iron chelator yielded remarkable synergy, significantly reducing the minimal inhibitory concentration. These results led to the hypothesis that other thiopeptides could also inhibit growth by using siderophore receptors to gain access to the cell. Here, we screened six thiopeptides for synergy with the iron chelator deferasirox against *P. aeruginosa* and a mutant lacking the pyoverdine receptors FpvA and FpvB. Our findings suggest that thiopeptides such as thiocillin cross the outer membrane using FoxA, the ferrioxamine siderophore receptor. Other structurally related thiopeptides did not inhibit growth of *P. aeruginosa*, but had greater potency against methicillin-resistant *Staphylococcus aureus* than thiostrepton and related thiopeptides. These results suggest that thiopeptide structures have evolved with considerations for target affinity and entry into cells.

---

Thiopeptides are a class of natural products with potent antibacterial activity against Gram-positive bacteria, including methicillin-resistant *Staphylococcus aureus* (MRSA)^1^. They are characterized chemically by the oxidation state of a central pyridine ring and by the size of a core macrocyclic ring comprised of thiazole moieties^2^. Thiopeptides act on the ribosome to inhibit protein translation but the exact mechanism depends on the number of members in the core macrocyclic ring: 26-membered macrocycles inhibit elongation factor G whereas 29-membered macrocycles block elongation factor Tu binding^3–5^. The exact mechanism of action for 35-membered macrocycles remains unknown, although they are reported to target the 50S ribosome, similar to thiostrepton (TS)^6^. TS is among the best-studied 26-membered thiopeptides and has multiple biological properties, including anticancer and antimalarial activities^7,8^.

In 1948, Su isolated a strain of *Micrococcus* from sewage samples that produced a compound with activity against Gram positive but not Gram negative bacteria^9^. The compound, micrococcin, was the first thiopeptide to be discovered. Since then, the thiopeptide family has expanded to over 100 members of diverse structure and origin^10^. Thiopeptides belong to a larger class of natural products called ribosomally synthesized and post-translationally modified peptides (RiPPs)^11^. These natural products are produced from a peptide precursor whose sequence varies depending on the thiopeptide, and undergo many post-translational modifications to reach their final structure^12^. Most thiopeptides are produced by Gram-positive soil and marine organisms including *Bacillus cereus, Streptomyces* spp., and *Nocardopsis* spp., although some Gram negatives such as *Serratia marcesens* have biosynthetic gene clusters encoding thiopeptide synthesis^13–17^.

Due to their poor solubility, challenging synthesis, and limited potency against Gram-negative bacteria, pharmaceutical companies ruled out thiopeptides for development as clinically useful antibiotics. Instead, researchers focused on addressing these issues through total synthesis, discovery of new thiopeptides and structure elucidation, identification of thiopeptide biosynthetic gene clusters, and modification of existing thiopeptides to improve solubility and activity ^12,17–25^. However, there is a gap in knowledge regarding their utility against Gram-negative bacteria which likely stems from the assumption that thiopeptides cannot cross the outer membrane.

We showed recently that TS hijacks pyoverdine receptors to cross the outer membranes of *Pseudomonas aeruginosa* and *Acinetobacter baumannii,* among the World Health Organization’s priority pathogens for new antibiotic development^26^ ^27^ *P. aeruginosa* and *A. baumannii* are opportunistic pathogens with intrinsic resistance to many antibiotics and are causative agents of healthcare-acquired infections^28^. Both express receptors for pyoverdine, a siderophore with high affinity for iron. TS uptake through pyoverdine receptors occurred under iron-limited conditions, and deferasirox (DSX), an FDA-approved oral iron chelator, strongly synergized with TS to inhibit the growth of clinical isolates of both species. Combinations of TS with other iron chelators were also effective^29^. These findings showed that thiopeptides can cross the outer membranes of specific Gram-negative bacteria. TS lacks activity against *Escherichia coli* because it does not express pyoverdine receptors. The thiopeptide family has over 100 members and recent bioinformatic analyses suggest that there could be hundreds more thiopeptides^30,31^. We hypothesized that other thiopeptides may also have activity against Gram-negative pathogens. Here, we screened six thiopeptides (Figure 1, Supplementary Figure 1) for activity against *P. aeruginosa* and potential synergy with DSX. Combinatorial assays showed that two of these used the ferrioxamine siderophore receptor, FoxA, to enter the cell, confirming our hypothesis that other thiopeptides could cross the outer membrane. The 26-membered macrocycles are of particular interest because modifications to the core macrocyclic ring altered siderophore receptor specificity. Other modifications abolished Gram-negative activity but improved Gram-positive activity. Thiopeptides are an underutilitzed class of antibiotics with potential for development as clinically useful drugs, especially as iron limitation at sites of infection could promote thiopeptide uptake.

**Figure 1.**
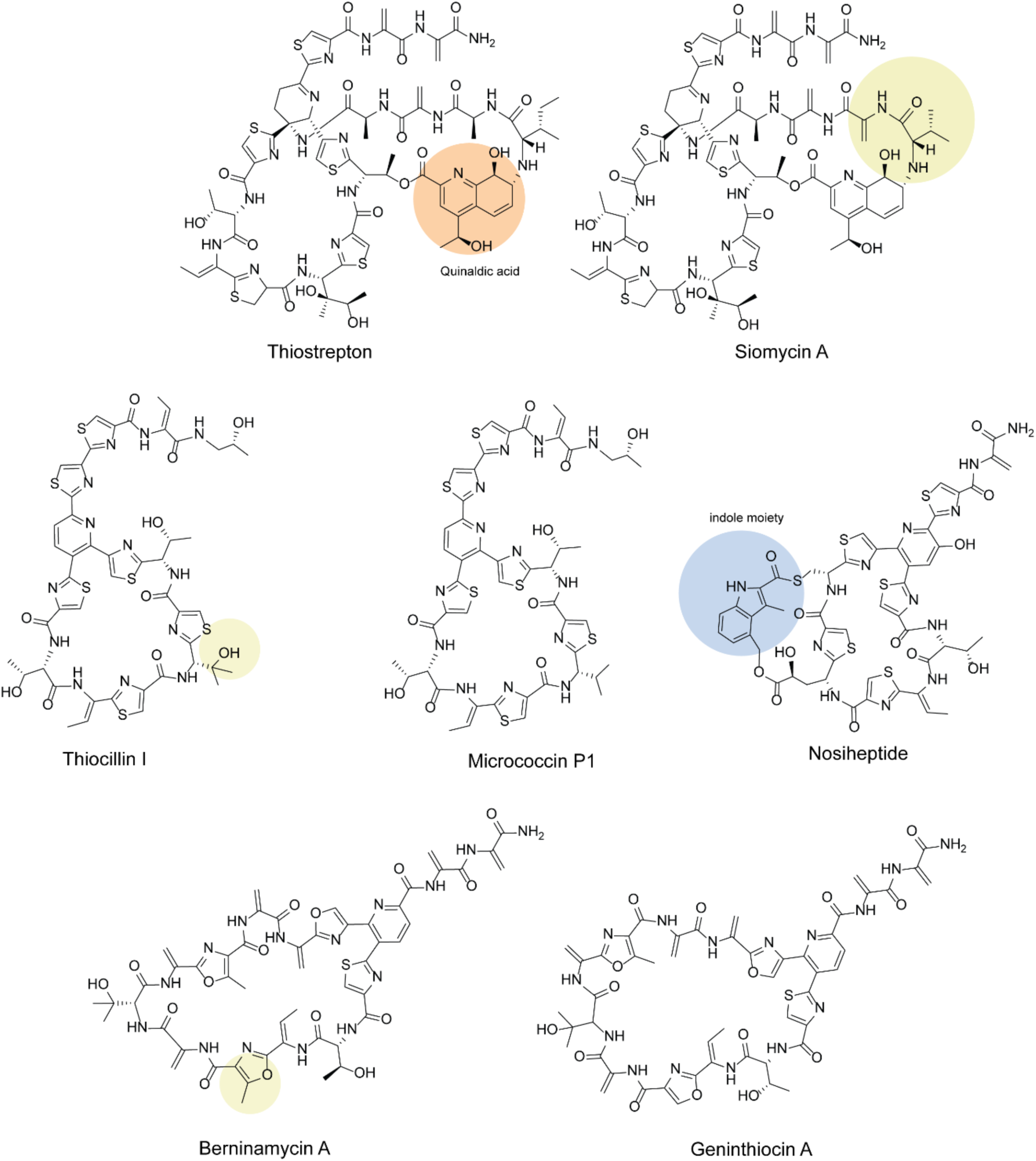
Structures of Thiopeptides Tested for Synergy with DSX. Differences between structural analogues are highlighted in yellow. The following are structural analogues: TS and SM, TC and MC, (26-membered macrocycles) and BER & GEN (35-membered macrocycles). NH is structurally similar to TC and MC, but has a unique indole moiety (blue). SM and TS have a unique quinaldic acid moiety (orange), but SM differs by two amino acids.

## RESULTS

### Siomycin A Uses the Pyoverdine Receptors for Uptake

We screened the activity of six thiopeptides in combination with DSX against *P. aeruginosa* PA14 using checkerboard assays. We looked for individual thiopeptide activity, synergy, and activity against a mutant lacking pyoverdine receptors FpvA and FpvB, which results in TS resistance. If the combination was active against the wild type but not the *fpvAB* mutant, the thiopeptide likely uses those receptors to cross the membrane. The six thiopeptides in the screen included siomycin A (SM), thiocillin I (TC), micrococcin P1 (MC), nosiheptide (NH), berninamycin A (BER), and geninthiocin A (GEN) (Figure 1).

SM differs from TS in that the two amino acids adjacent to the quinaldic acid moiety are dehydroalanine and isoleucine instead of alanine and leucine. TC and MC are nearly identical except that TC has a hydroxyvaline and MC has a valine in its macrocyclic ring. BER and GEN differ by a single methyl group on one of the oxazole rings. NH resembles TC and MC but is structurally unique from the other thiopeptides, with an indole moiety linked to the core macrocyclic ring to form a secondary ring. TS, SM, TC, MP, and NH belong to the 26-membered macrocycles whereas BER and GEN have 35-membered macrocycles.

Based on checkerboard assays, BER, GEN, and NH did not synergize with DSX (Supplementary Figure 1). SM synergized with DSX against wild type PA14 but the combination had no effect against the *fpvAB* mutant (Figure 2). Unlike the wild type, growth of the mutant was inhibited by 16 µg/ml DSX alone. These data are similar to those obtained previously with TS + DSX^26^ and indicates that SM likely crosses the outer membrane using the pyoverdine receptors. However, the synergistic profile of SM + DSX was less pronounced compared to TS + DSX. There are two potential explanations for its weaker synergistic profile: 1) siomycin has less affinity for the target or 2) siomycin has less affinity for the receptors. To distinguish between these possibilities, we tested each thiopeptdide against the Gram-positive pathogen MRSA USA 300 in a broth dilution assay (Supplementary Figure 2). MRSA lacks the outer membrane; therefore, the main consideration would be differences in affinity for the highly-conserved ribosomal target. No differences between the minimal inhibitory concentration (MIC) of cells treated with TS or SM were observed, suggesting that the differences in susceptibility in *P. aeruginosa* were related to receptor affinity. Further, this result provides some structure-activity relationship (SAR) information regarding interactions between TS and the pyoverdine receptors, specifically that the alanine and leucine adjacent to the quinaldic acid do not affect target activity but improve uptake; however, they are not essential because SM can still use the pyoverdine receptors to cross the outer membrane.

**Figure 2.**
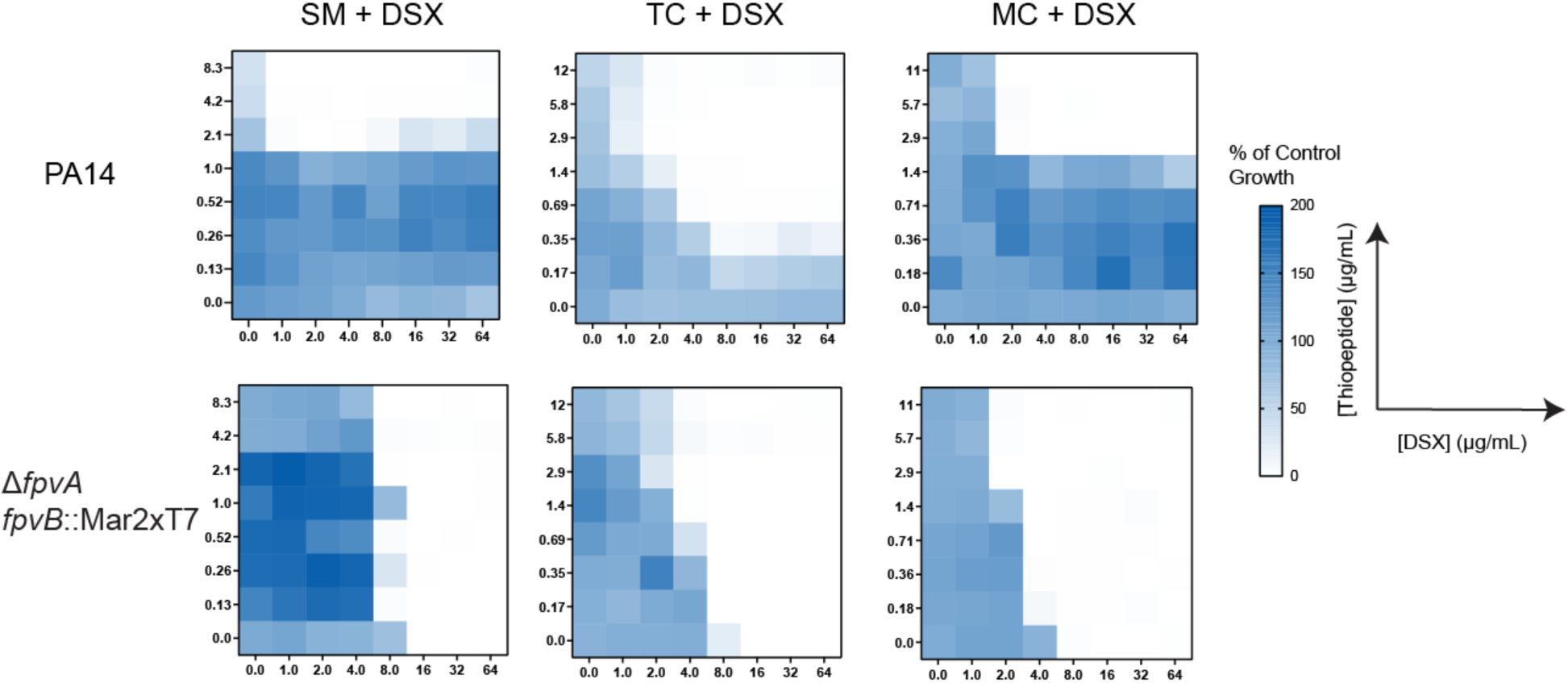
Thiopeptides synergize with DSX against PA14 and PA14 *ΔfpvA fpvB*::Mar2xT7. Checkerboard assays were conducted in 10:90 medium. Top row are checkerboards conducted with each thiopeptide against wild type PA14 whereas bottom row are checkerboards conducted against its pyoverdine receptor-null mutant. The concentration of the thiopeptides are indicated along the column (maximum concentration of 10 µM in µg/mL except for SM which was tested at 5 µM) and the concentration of DSX is indicated along the row (maximum concentration of 64 µg/mL). Growth is proportional to the intensity of blue whereas white indicates no growth. Results are averaged from three independent experiments.

### Thiocillin I and Micrococcin P1 Exploit the Ferrioxamine Receptor

In checkerboard assays, TC and MC synergized with DSX against *P. aeruginosa* PA14 (Figure 2). The concentration of DSX was kept constant at 64 µg/mL, while the highest concentration of TC and MC tested due to solubility limitations was 12 µg/mL (10 µM). Remarkably, this combination inhibited growth of the *fpvAB* mutant (Figure 2). This result indicated that that TC and MC likely use a different siderophore receptor to enter the cell. To identify that receptor, we screened a panel of 16 siderophore receptor-deficient transposon mutants of PA14 (Supplementary Table 2). Only a *foxA* transposon mutant, lacking the ferrioxamine receptor, displayed resistance to TC + DSX (Figure 3AB). When the mutant was complemented with *foxA* in trans, susceptibility to TC was restored, confirming that *foxA* is necessary for TC uptake (Figure 3C). The susceptibility of the complemented strain was slightly reduced compared to the wild type, likely due to the low copy number of the vector in *P. aeruginosa*.

**Figure 3.**
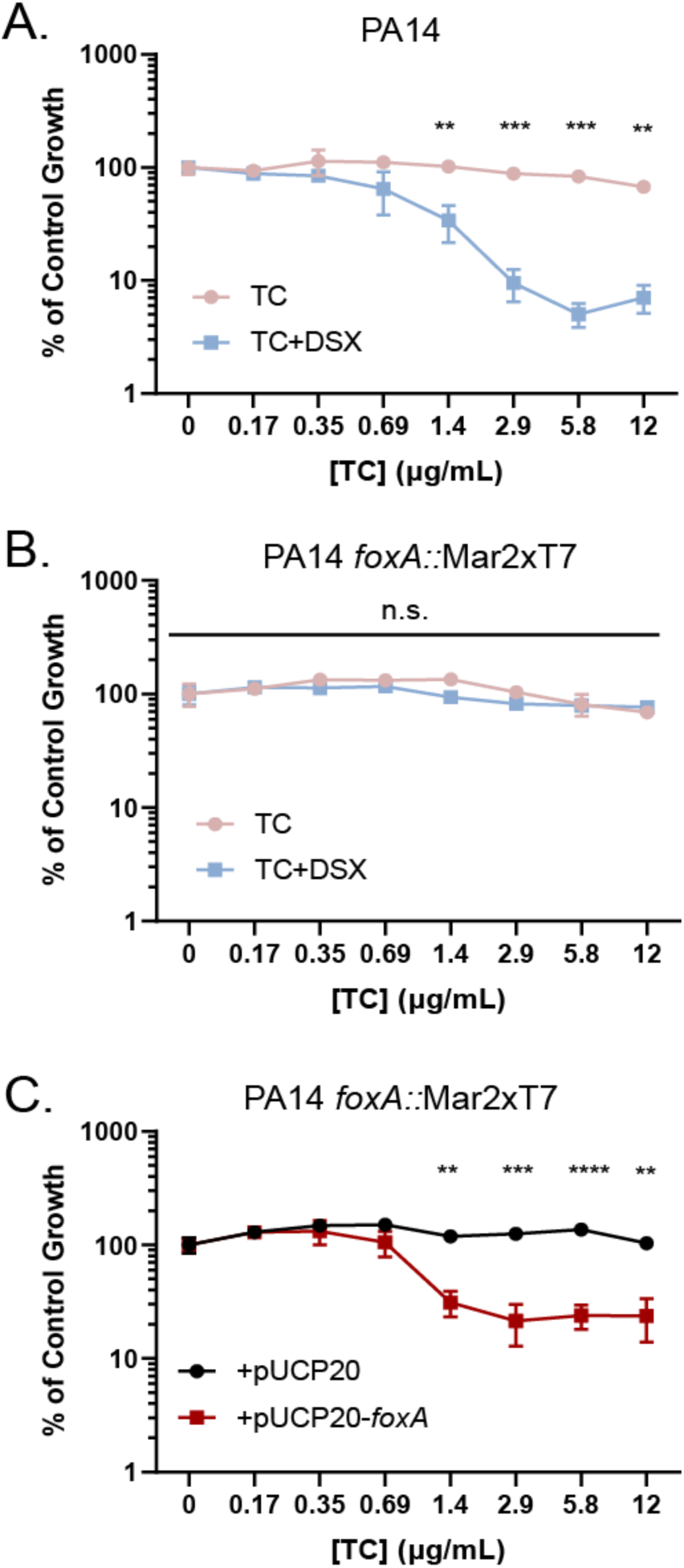
TC crosses the outer membrane through the FoxA siderophore receptor. Wild type PA14 (A) and PA14 foxA::Mar2xT7 (B) were challenged with TC (pink) and TC + DSX (blue). Dose response assays were conducted in 10:90. The transposon mutant was complemented with wild type PA14 foxA (C) and challenged with TC in VBMM. The vector control is in black whereas the complemented mutant is in red. Results were averaged from three independent replicates. n.s, not significant; **, p<0.01; ***, p<0.0005; ****, p<0.0001. Significance was determined using two-way ANOVA followed by Sidak’s multiple-comparison test.

### TC and MC inhibit the growth of TS-resistant clinical isolates

In our previous studies, we identified *P. aeruginosa* clinical isolates resistant to the TS + DSX combination. One isolate, C0379, was resistant due to a 5’ truncation of the gene encoding the FpvB receptor^26,29^. We hypothesized that TC or MC might inhibit the growth of this isolate because the only differences in its *foxA* gene compared to that of PAO1 were five silent mutations and a single base pair mutation (D760G) in one beta-strand belonging to a structural component of the receptor (Supplementary Table 3). Consistent with this idea, C0379 was susceptible to combinations of TC + DSX or MC + DSX (Supplementary Figure 3A). Bioinformatic analysis showed that *A. baumannii* also encodes the FoxA receptor. We tested one multidrug-resistant clinical isolate, C0286, for susceptibility to TC + DSX and found that it was susceptible even at the lowest concentration of TC tested (0.17 µg/mL) (Supplementary Figure 3B). However, the lab strain ATCC 17978 was not susceptible to the combination, suggesting inter-strain differences. This is not unusual as it is well established that lab adapted strains may not behave similarly to clinical isolates^32^. Overall, we conclude that TC + DSX can inhibit the growth of *P. aeruginosa* and *A. baumannii* clinical isolates including those resistant to TS.

### A cocktail of TS + TC + DSX has an additive effect

Our observation that thiopeptides synergize with DSX against *P. aeruginosa* demonstrated the potential to target gram-negatives expressing either pyoverdine or ferrioxamine receptors using thiopeptide-chelator cocktails. Sub-MIC combinations of TS (0.1 µg/mL) + 64 µg/mL DSX and TC (0.2 µg/mL) + 64 µg/mL DSX, as well as TS + TC + DSX were tested against PA14 in VBMM (Figure 4). Each double combination inhibited growth to ∼30% of control; however, the triple combination decreased the growth to only 10% of control. A multiplicative rule approach was used to determine the effect of combining both thiopeptides where the expected growth is the product of the growth inhibition for the individual combinations (i.e. TS + DSX or TC + DSX). For TS + DSX, growth was reduced to 31% of control and for TC + DSX, growth was reduced to 29% of control. Therefore, the expected growth for the combination of TC + TS + DSX would be 9% of control. DSX alone had no effect on growth. The experimental value yielded 8.8% of control, suggesting an additive effect of the thiopeptides. These results indicate that thiopeptide cocktails could offer an advantage compared over single thiopeptide treatment by using different siderophore receptors to enter the cell.

**Figure 4.**
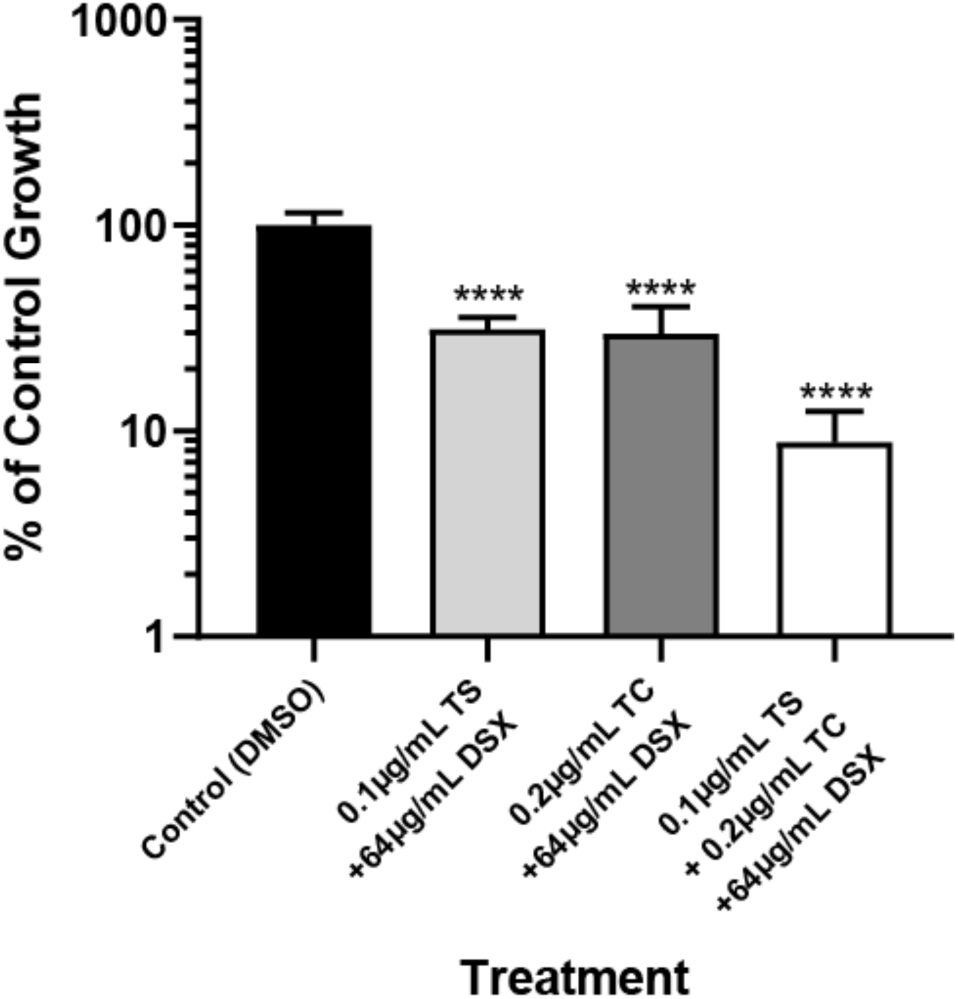
A thiopeptide cocktail significantly reduces PA14 growth. Control growth less than 20% was considered as the MIC cut-off. Assays were conducted in VBMM. Results are averaged from three independent experiments. ****,p<0.0001. Significance was determined using one-way ANOVA followed by Tukey honest significant difference post test.

### Thiopeptides target specific bacterial genera and species

Unlike the pyoverdine receptors, the ferrioxamine receptor is not unique to Pseudomonads. Therefore, we tested whether other Gram-negative bacteria were susceptible to TC. Bioinformatic analyses of other priority Gram-negative pathogens such as *Escherichia coli, Klebsiella pneumoniae,* and *Salmonella* Typhimurium showed that they carry *foxA* in their genome. However, they were not susceptible to TC or the combination of TC + DSX at the highest concentrations tested, suggesting that the receptors are sufficiently different in sequence to prevent recognition of TC as a ligand. FoxA proteins from *S.* Typhimurium SL1344 and *P. aeruginosa* PAO1 share 40% amino acid sequence identity (Supplementary Figure 4). Residues Y218, H374, and Q441 that interact with the native ferroxamine ligand are conserved in both species, indicating that TC and MC must rely on interactions with other amino acids for uptake into Pseudomonads^33^.

To determine if there were differences in susceptibilities between different species of *Pseudomonas*, we conducted dose-response assays with TC and TC + DSX against *P. putida* KT2440, *P. marginalis* CVCOB 1152, *P. protegens* Pf-5, and *P. fluorescens* PV5(Supplementary Figure 5. Similar experiments were done with TS and TS + DSX. We defined resistance if the percent of control growth exceeded 80%, susceptible if below 20%, and intermediate if between 20-80%. Interestingly, there was variability in the susceptibility to the combinations between species. *P. putida* was resistant to TS but susceptible to TC (Figure 5, Supplementary Figure 5). *P. protegens* was resistant to all combinations (Figure 5, Supplementary Figure 5). Alignments of the FoxA sequences for each species showed 65%, 65%, and 60% similarity for *P. fluorescens, P. protegens,* and *P. putida* compared to *P. aeruginosa* PAO1. Despite the differences in susceptibility to TC + DSX, the key residues that interact with the native ligand are conserved, similar to *S.* Typhimurium, suggesting that TC interacts with other amino acids. To focus on possible sites of interaction, we looked for residues shared between *P. aeruginosa, P. putida*, and *P. fluorescens*, but not *P. protegens*, which was resistant to TC and TC + DSX (Figure 5, Supplementary Figure 5). A total of 17 residues were identified following this criterion and mapped onto a co-crystal structure of FoxA bound to ferrioxamine (PDB ID: 6I96)^33^. Only three were found in proximity to the native ligand – A620(P), S612(A), and D380(N) (Supplementary Table 4). A629(P) indicates that in the FoxA receptor of *P. aeruginosa, P. putida,* and *P. fluorescens* at position 620, the amino acid is alanine. However, in *P. protegens* the amino acid at the same position is proline. A620(P) and S612(A) are located in loop 7 of FoxA, which closes upon interaction of the ligand with the active site^33^. Closure of loop 7 prevents the native ligand from dissociation. D380(N) is close to conserved residue H374, which interacts with the native ligand.

**Figure 5.**
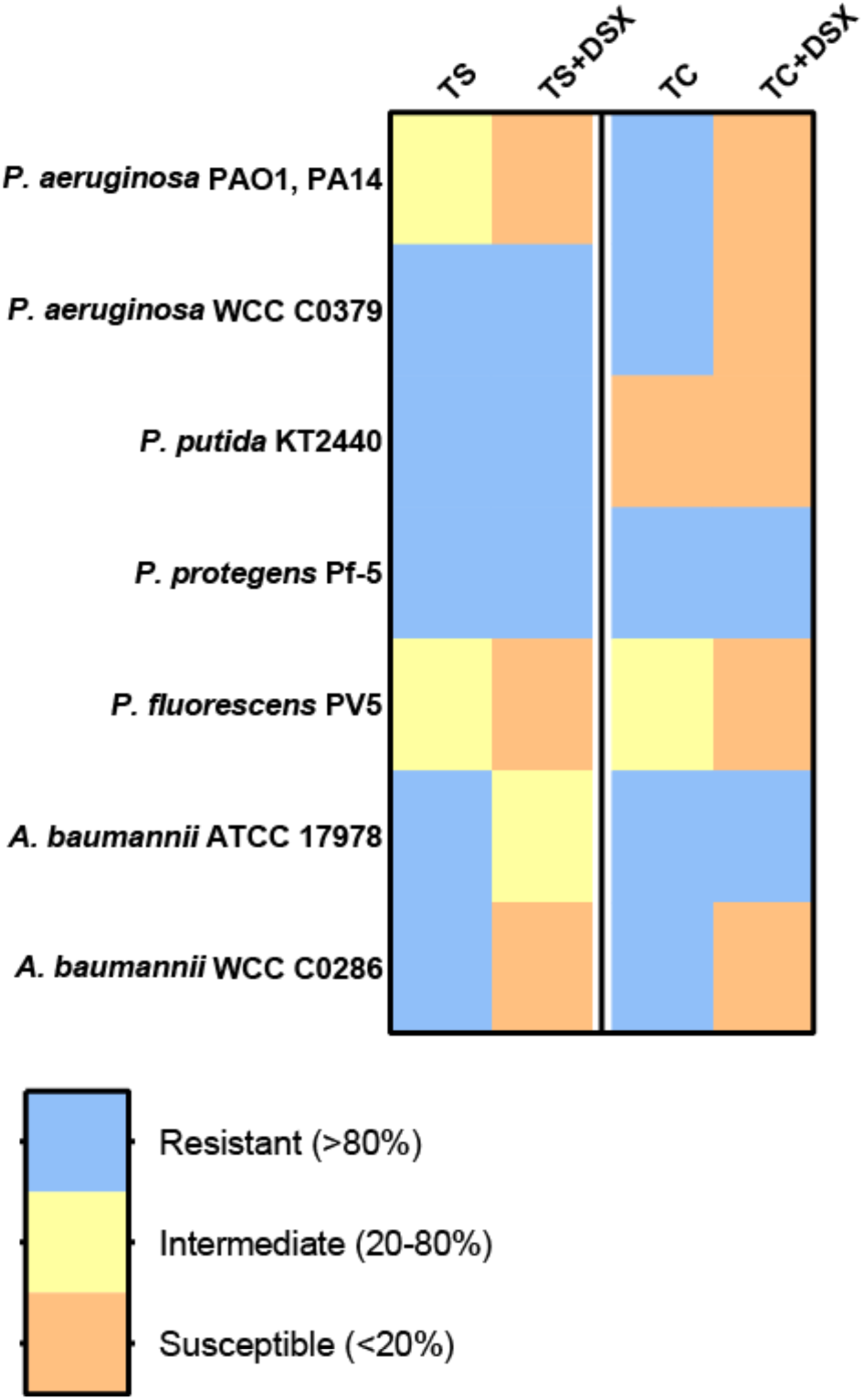
Susceptibility to TS and TC are different between species and strains. Susceptibility of *Pseudomonas* and *Acinetobacter* to TS and TC in combination with DSX. Blue boxes indicate resistance (>80% of control growth). Yellow boxes indicate intermedicate susceptibility (20-80% of control growth). Orange boxes indicate susceptible (<20% of control growth).

A BLAST search using the PAO1 FoxA amino acid sequence against *P. aeruginosa, P. putida, P. fluorescens* (all TC susceptible under low iron), and *P. protegens* (TC resistant) was conducted to see if A620(P), S612(A), and D380(N) are conserved (Supplementary Figure 6). The top 100 hits were aligned (MUSCLE), which revealed that alanine at position 620 and aspartic acid at position 380 are well conserved in *P. aeruginosa, P. fluorescens,* and *P. putida*. Alanine was the most common amino acid at position 612 in *P. fluorescens* rather than serine, suggesting that this residue may not be important for TC interaction with FoxA since this species was susceptible to TC + DSX. In *P. protegens,* which is resistant to TC + DSX, lysine was the most abundant at residue 620. At residue 380, glutamine occurred more frequently. These differences suggest that alanine and aspartic acid at residues 620 and 380 respectively may be involved in TC uptake. These data suggest that TC uptake requires specific amino acid interactions that differ between species.

### Nosiheptide is a potent inhibitor of MRSA USA 300

The core structure of NH is nearly identical to TC and MC; however, a single indole is linked to the core 26-membered macrocyclic ring (Figure 1). This small change in structure results in loss of synergy with DSX (Supplementary Figure 1). Although synergy was not observed, there was a reduction in growth when NH was combined with DSX. When tested against the *fpvAB* mutant, potentiation was lost (Supplementary Figure 1).

There are examples – such as psychotrimine and daptomycin – of natural product antibiotics containing tryptophan or its side chain, indole ^34^. Daptomycin is a Gram-positive antibiotic that disrupts the cytoplasmic membrane^35^. Like NH, daptomycin is a bulky molecule whose size prevents it from crossing the outer membrane, making it ineffective against Gram-negative bacteria. Based on its unique indole content, we hypothesized that NH might have improved efficacy against Gram-positive bacteria compared to the other thiopeptides tested. All seven thiopeptides were tested in a dose-response assay against MRSA USA 300, with 260 ng/mL as the highest concentration (Supplementary Figure 2). Consistent with our hypothesis, NH was the most potent against MRSA, with a MIC of <4 ng/mL, followed by TS and SM (32 ng/mL), TC and MC (130 ng/mL), and BER and GEN (>260 ng/mL). These results show that the indole moiety of NH may confer an advantage against Gram-positives compared to other thiopeptides.

## DISCUSSION

We previously showed that TS hijacked the pyoverdine receptors of *P. aeruginosa* and *A. baumannii* to cross the outer membrane. These data informed the hypothesis that other thiopeptides might similarly act as Trojan Horse antibiotics. In this work, we explored the ability of six other thiopeptides to exploit siderophore receptors under iron-limited conditions. We showed that SM – a thiopeptide structurally similar to TS – also uses pyoverdine receptors to enter cells. The differences in structure of the two compounds provided SAR information. The IC_50_ of TS and SM are similar (0.1 µM vs 0.25 µM) based on *in vitro* studies of *E. coli* ribosomal activity^36^. This suggests that SM has weaker synergy with DSX because of reduced affinity for the receptor compared to TS. Combinatorial analyses by checkerboard assays suggested that TC and MC used a different siderophore receptor. We screened mutants lacking various receptors in low iron growth media, leading to the discovery that those thiopeptides used FoxA, a xenosiderophore receptor in *P. aeruginosa*. TC was also effective against a TS-resistant *P. aeruginosa* clinical isolate. However, despite widespread expression of FoxA homologues, susceptibility to TC was not universal among Gram-negative bacteria. Even within the genus *Pseudomonas*, there was variability in susceptibility between species. Differences in the levels of siderophore receptor expression could be a factor in the susceptibility profiles as a higher abundance would allow more thiopeptide uptake.

This work suggests links between thiopeptide structure and activity against specific Gram-negatives. Thiopeptides were thought to have little or no activity against Gram-negative pathogens due to their inability to cross the outer membrane. However, thiopeptide producers are mainly marine and soil bacteria, which inhabit environments shared by Pseudomonads. We propose that thiopeptide structures evolved to target the siderophore receptors of cohabiting Gram-negatives. This is not unprecedented as many other natural products can use such receptors to cross the outer membrane. For example, *P. aeruginosa* produces pyocins that target the pyoverdine receptors of competing strains^37^.. NH failed to synergize with DSX, although the combination reduced growth. Potentiation was lost against the *fpvAB* mutant. This result suggests that NH could have poor affinity for pyoverdine receptors, but further work needs to be done to demonstrate this. We also showed that of the thiopeptides tested, NH had the greatest activity against MRSA USA 300. The exact mechanism remains unknown, but while the core of NH is nearly identical to TC and MC, the indole moiety makes it unique compared to the other thiopeptides. NH does not synergize with DSX which suggests that the indole moiety could interfere with its recognition by the FoxA receptor. Tryptophan and indole increase interactions of peptides with membranes and can act as signaling molecules^38–41^. We speculate that the indole moiety may provide enhanced peptide-membrane interactions, facilitating the ability of NH to cross the membrane and target Gram-positive pathogens like MRSA. We observed a lower MIC with NH (<4 ng/mL) compared to TS (32 ng/mL) against MRSA in 10:90 medium, consistent with results from other groups using Mueller-Hinton Broth^1,42^. In contrast, assays using purified ribosomes showed that TS had 40% greater inhibitory activity compared to NH^43^. While NH has weaker inhibitory activity against purified ribosomes, it may compensate with improved membrane permeability. NH has been tested against MRSA in an intraperitoneal mouse infection model^1^. Mice treated with 20 mg/kg NH showed 90% survival over 5 days compared to 40% in the vehicle control group.

Comparing TC and TS, it is apparent that the core macrocycle is nearly identical. However, TS has a second macrocyclic ring with a structurally-distinct quinaldic acid moiety. It is likely that this second ring allows TS to interact with the pyoverdine receptors rather than FoxA. In contrast, BER and GEN had no activity against *P. aeruginosa* under iron-limitation. Even against MRSA, these 35-membered macrocycle thiopeptides had no activity at the highest concentration tested (260 ng/mL). This is unsurprising as it was previously reported that BER has weaker inhibitory activity on purified ribosomes compared to TS^6^. Our results do not rule out the potential of 35-membered thiopeptides having other advantages. Furthermore, in this study we did not test 29-membered thiopeptides such as GE2270A, thiomuracin A, and GE37468^44–46^. These molecules could also have potential activity against *P. aeruginosa* and *A. baumannii* under iron-limited conditions.

Schwalen et al. recently used a bioinformatics approach to search for enzymes that catalyze non-spontaneous [4+2]-cycloadditions characteristic of thiopeptide biosynthesis. They identified 508 [4+2]-cycloaddition enzymes, 372 of which were unique at the amino acid level^31^. These results suggest that there are many more structurally unique thiopeptides than previously estimated. Diverse thiopeptides may reveal interesting new activities with antimicrobial applications. One example is cyclothioazomycin, a structurally distinct compound with activity against *Bacillus* spp., although weaker relative to other thiopeptides^47^. Interestingly, this thiopeptide has potent antifungal activity via its interactions with chitin^48^. It also was reported to be a human plasma renin inhibitor^49^.

Trojan horse antibiotics are an interesting avenue for drug development. They have advantages over antibiotics that must simply diffuse through the outer membrane because of their greater specificity and uptake. Moreover, sites of infection are frequently iron-limited, which naturally potentiates such molecules. Hosts produce iron sequestration proteins such as transferrin and lactoferrin that restrict iron, a process called nutritional immunity^50–52^.

While thiopeptides been overlooked for use in the clinic, we show the potential for these antibiotics to be developed as selective Gram-negative antimicrobials that can target high priority pathogens including *P. aeruginosa* and *A. baumannii*. Our work also supports the future development of NH as an anti-MRSA therapeutic.

## METHODS

### Chemicals

Thiopeptides were purchased from Cayman Chemicals and stored at −20°C. Thiopeptide purity was at least ≥95%. All thiopeptides were dissolved in dimethylsulfoxide (DMSO) with no solubility issues ranging from 1mg/mL to 20mg/mL. Deferasirox was purchased from AK Scientific (98% purity). A stock solution of 20 mg/mL in DMSO was made. Stock solutions were stored at −20°C until use and all serial dilutions were prepared using fresh DMSO at 75x the final concentration.

### Bacterial Strains and Culture Conditions

All bacterial strains used in this study are listed in Supplementary Table 1. Bacteria were inoculated from glycerol stocks stored at −80°C and grown overnight in LB at 37°C with shaking (200 rpm). Bacteria were subcultured (1/500 dilution) into fresh 10:90 media or Vogel-Bonner Minimal Media (VBMM) and grown for 16 hours. Optical densities were standardized to OD_600_ 0.1 and diluted 1/500 into fresh 10:90 or VBMM, which was used in microbroth dilution MIC assays or checkerboard assays.

### Generation of PA14 f*oxA*::Mar2xT7 pUCP20-*foxA*

*foxA* was PCR amplified from wild type PA14 genomic DNA prepared using Qiagen genomic DNA isolation kit. The following forward and reverse primers were used to obtain wild type *foxA* with SacI and Xbal restriction sites respectively (underlined): forward 5’GAAG<u>GAGCTC</u>GTGGACGCTTGCTTTCGT and reverse 5’GCGT<u>TCTAGA</u>GACGCCGGCGAATGCCC. The PCR product and pUCP20 were digested with SacI and Xbal (Thermo Scientific) using Thermo Scientific Fast Digest. The products were run on a 1% agarose gel for 30mins at 120V, cut out, and purified using GeneJET Extraction Kit (Thermo Scientific). The purified linearized vector and digested *foxA* gene were ligated (1:3 molar ratio) overnight at 4°C using T4 DNA ligase (Thermo Scientific). The construct was transformed into competent *E. coli* DH5a by heat shock at 42°C and plated on LB plates with 100 µg/mL ampicillin and 5-bromo-4-chloro-3-indolyl-β-D-galactopyranoside (IPTG) for blue-white screening (BioShop). Plates were incubated at 37°C overnight. White single colonies were streaked on fresh LB plates with ampicillin to confirm that colonies contained the construct. Minipreps were done using GeneJET Plasmid Miniprep Kit (Thermo Scientific) and quantified by NanoDrop. The *foxA* transposon mutant was grown overnight in LB. 1 mL was centrifuged and washed twice with nuclease-free water (Qiagen). The cell pellet was resuspended in 500 µL of nuclease-free water. 100 ng of the construct was added to 100 µL of resuspended cells and electroporated (2.50 V). After a recovery period of 2 hours, cells were plated onto LB agar with 200 µg/mL carbenicillin (Bioshop). Plates were incubated at 37°C overnight. Single colonies were streaked on fresh plates to confirm successful electroporation. A single colony was picked and grown in liquid for MIC assays. The susceptibility of the *foxA* transposon mutant harbouring pUCP20-*foxA* to TC indicated successful complementation. Similarly, the same *foxA* transposon mutant was electroporated with empty pUCP20 as vector control.

### Microbroth Dilution and Checkerboard Assays

Stock solutions were diluted in DMSO and 2µL of the resulting solutions plus 148µL of a bacterial suspension standardized to an OD600 of ∼ 0.1 and diluted 1:500 in 10:90 was added to a 96 well plate (Nunc) in triplicate. Control wells contained 148µL of 10:90 + 2µL DMSO (sterility control) or standardized bacterial suspension + 2µL DMSO (growth control). After incubating at 37°C for 16h, with shaking at 200 rpm, the OD600 was read using a plate reader (Thermo Scientific). Assays were performed in triplicate, calculations were done in Microsoft Excel, and results were graphed using Prism (GraphPad) as a percentage of the DMSO control. For wells with 2 compounds, such as increasing TS concentrations with 64µg/mL of DSX, 4µL of DMSO was used in the sterility and growth controls.

Checkerboard assays were set-up using Nunc 96-well plates in an 8-well by 8-well format. Two columns were allocated for vehicle controls and two columns for sterility controls. Vehicle controls contained 4 µL DMSO, in 146 µL of 10:90 inoculated with PA14 as described in Growth Curves. Sterile controls contained the same components in 10:90, without cells. Serial dilutions of each thiopeptide – with 10 µM being the highest final concentration – was added along the ordinate of the checkerboard (increasing concentration from bottom to top) whereas serial dilutions of DSX – with 64 µg/mL being the highest final concentration – was added along the abscissa (increasing concentration from left to right). The final volume of each well was 150 µL and each checkerboard was repeated at least three times. Plates were incubated and the final OD600 determined as detailed above. Checkerboards were graphed as heatmaps in Prism (GraphPad).

### Bioinformatic Analysis

MUSCLE alignments were conducted using Unipro UGENE and Geneious Prime. Accession numbers of *P. aeruginosa, P. fluorescens, P. protegens, P. putida,* and *S.* Typimurium FoxA are as follows: AAG05854.1, WP_122305244.1, WP_003218248.1, AAY95537.2, NP_742329.1, and CBW16459.1. Genomic analyses for mutations in WCC C0379 were done with breseq by Kara Tsang using *P. aeruginosa* PAO1 as the reference genome.

## ASSOCIATED CONTENT

### Supporting Information

Table S1, list of strains; Table S2, mutants tested to show that TC uses the FoxA receptor to cross the outer membrane; Table S3, breseq genomic annotations of the FoxA receptor for WCC C0379; Table S4, FoxA amino acid residues that are different in *P. protegens* compared to other *Pseudomonas* spp; Figure S1, checkerboard assays for thiopeptides that did not synergize with DSX; Figure S2, thiopeptide activity against MRSA USA300; Figure S3, thiopeptide activity against *P. aeruginosa* and *A. baumannii* TS-resistant clinical isolates; Figure S4, MUSCLE alignment of *S.* Typhimurium and *P. aeruginosa* FoxA receptors; Figure S5, dose-response assays of thiopeptides against different *Pseudomonas* spp. Figure S6, Sequence logos of FoxA amino acid sequences of *Pseudomonas* spp.

## Author Contributions

DCKC performed all experiments. DCKC and LLB wrote the paper.

## Funding Sources

This work was funded by grants to LLB from the Natural Sciences and Engineering Research Council (NSERC) RGPIN-2016-06521 and the Ontario Research Fund RE07-048. DCKC holds an NSERC Canadian Graduate Scholarship – Master’s Award.

## ACKNOWLEDGMENTS

We thank Gerry Wright for strains from the Wright Clinical Collection, as well as Eric Brown and Brian Coombes for the following strains: MRSA USA 300 and *K. pneumoniae*, and S. Typhimurium SL1344. We thank John Whitney for the following strains: *P. protegens* Pf-5, and *P. putida* KT2440 and David Guttman for *P. fluorescens*. We thank Kara Tsang for her help with breseq analyses. We thank Christy Groves for assistance with the abstract graphic.

## ABBREVIATIONS

TS,: thiostrepton;
TC,: thiocillin;
MC,: micrococcin;
NH,: nosiheptide;
MIC,: minimal inhibitory concentration;
SM,: siomycin;
BER,: berninamycin;
GEN,: geninthiocin;
DSX,: deferasirox;
FpvA,: ferripyoverdine receptor A;
FpvB,: ferripyoverdine receptor B;
FoxA,: ferrioxamine receptor;
WCC,: Wright Clinical Collection;
VBMM,: Vogel-Bonner Minimal Media;
MRSA,: methicillin-resistant *Staphylococcus aureus*;
*E. coli*,: *Escherichia coli*;
*K. pneumoniae*,: *Klebsiella pneumoniae*;
*S.* Typhimurium,: *Salmonella* Typhimurium;
*P. aeruginosa,*: *Pseudomonas aeruginosa*;
*P. fluorescens*,: *Pseudomonas fluorescens*;
*P. putida*,: *Pseudomonas putida*;
*P. protegens,*: *Pseudomonas protegens;*
*A. baumannii*: *Acinetobacter baumannii*

---

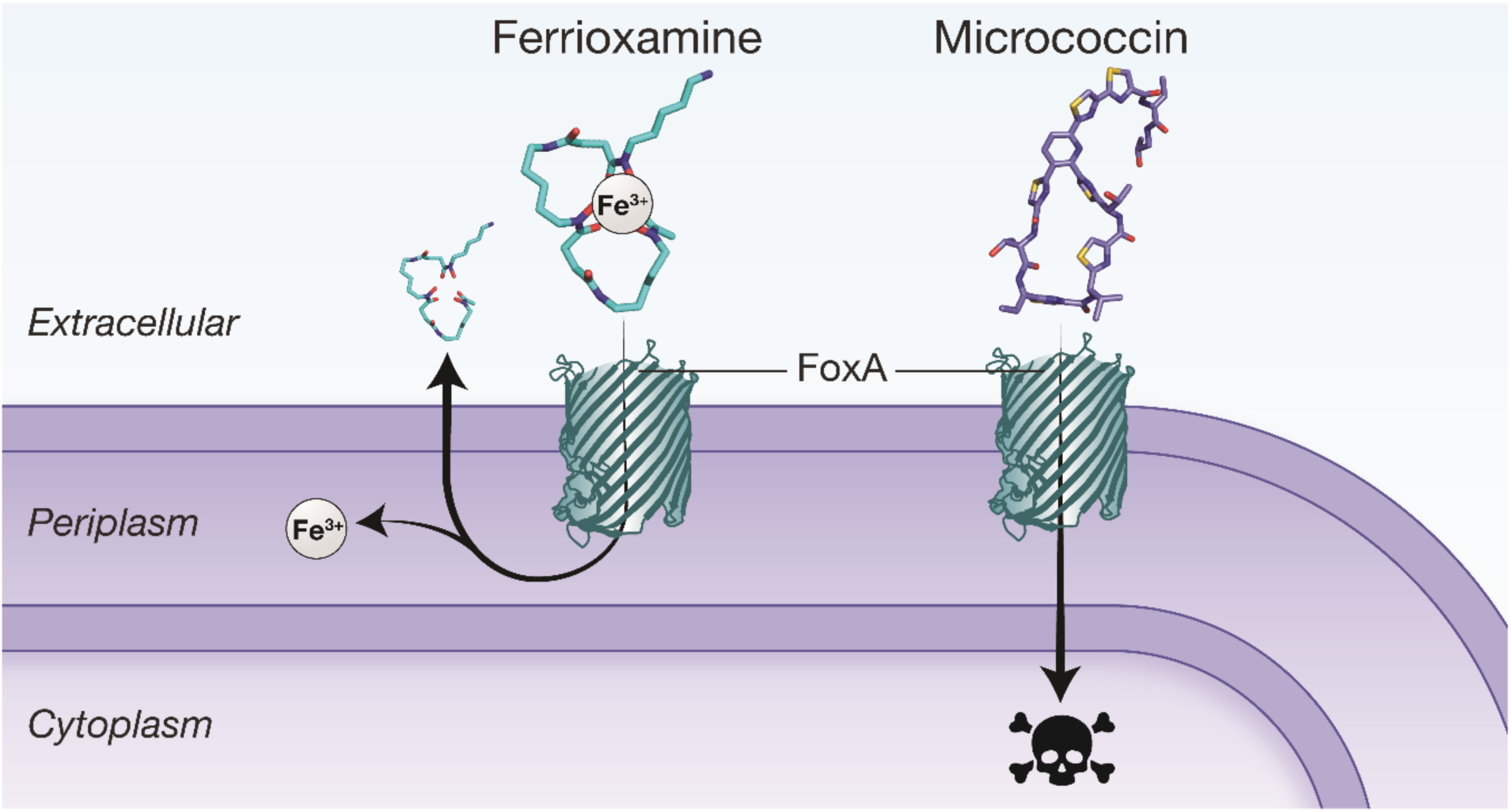
For Table of Contents Only

## Notes

### Competing Interest Statement

The authors have declared no competing interest.

### Summary of Updates

Corrected structure for thiostrepton.

