## Supplementary Tables S1-4, Figures S1-S6, references. for "Thiocillin and Micrococcin Exploit the Ferrioxamine Receptor of *Pseudomonas aeruginosa* for Uptake" (from another version of this preprint: version 1, 2020-04-23)

**Supplementary Table 1:** Strains used in this study.

| **Strain** | **Reference** |
| --- | --- |
| *P. aeruginosa* PA14 | ^1^ |
| *P. aeruginosa* PA14 *foxA*::Mar2xT7 | ^1^ |
| *P. aeruginosa* PA14 ∆*fpvA fpvB*::Mar2xT7 | ^2^ |
| *Pseudomonas fluorescens* PV5 | Guttman Lab |
| *Pseudomonas protegens* Pf-5 | Whitney Lab |
| *Pseudomonas putida* KT2440 | Whitney Lab |
| MRSA USA 300 | Brown lab |
| *Klebsiella pneumoniae* | Brown Lab |
| *Escherichia coli* K12 BW25113 | ^3^ |
| *E. coli* DH5ɑ | This study |
| *Salmonella* Typhimurium SL1344 | Coombes Lab |

**Supplementary Table 2:** Panel of Transposon Mutants Used to Screen for TC and MC resistance. The *foxA* transposon mutant is highlighted.

| **Strain/mutant** | **Gene(s)** | **Function** |
| --- | --- | --- |
| *fpvA*::Mar2xT7 | PA2398 | Type 1 ferripyoverdine receptor |
| *fpvB*::Mar2xT7 | PA4168 | Alternate Type 1 ferripyoverdine receptor |
| ∆*fpvA fpvB*::Mar2xT7 | PA2398, PA4168 | Type I ferripyoverdine receptor, Alternate type I ferripyoverdine receptor |
| *fptA*::Mar2xT7 | PA4221 | Pyochelin receptor |
| PA1322::Mar2xT7 | PA1322 | Probable TonB-dependent receptor |
| PA4837::Mar2xT7 | PA4837 | Probable outer membrane protein precursor |
| ***foxA*::Mar2xT7** | **PA2466** | **Ferrioxamine receptor FoxA** |
| *fiuA*::Mar2xT7 | PA0470 | Ferrichrome receptor FiuA |
| *pirA*::Mar2xT7 | PA0931 | Alternate enterobactin receptor |
| *pfeA*::Mar2xT7 | PA2688 | Enterobactin receptor |
| *hasR*::Mar2xT7 | PA3408 | Heme uptake outer membrane receptor |
| *fvbA*::Mar2xT7 | PA4516 | Vibriobactin receptor |
| *piuA*::Mar2xT7 | PA4514 | Hydroxamate-type ferrisiderophore receptor |
| PA0151::Mar2xT7 | PA0151 | Probable TonB-dependent receptor |
| *chtA*::Mar2xT7 | PA4675 | Aerobactin, Rhizobactin 1021, Schizokinen receptor |
| *phuR*::Mar2xT7 | PA4710 | Heme/Hemoglobin uptake receptor precursor |

**Supplementary Table 3.** WCC C0379 has mutations in the FoxA receptor that do not affect TC uptake. Analysis was done using breseq with PAO1 as the reference genome.

| **Position** | **Mutation** | **Type** |
| --- | --- | --- |
| 2,782,920 | G769G (GGT→GGC) | Silent |
| 2,782,948 | D760G (GAC→GGC) | Single base pair |
| 2,783,001 | A742A (GCT→GCG) | Silent |
| 2,784,231 | D332D (GAT→GAC) | Silent |
| 2,784,519 | G236G (GGT→GGC) | Silent |
| 2,785,134 | L31L (CTG→CTA) | Silent |

**Supplementary Table 4:** FoxA amino acid residues different in *P. protegens.* 17 amino acids in the FoxA receptor differ between *P. protegens* and *P. aeruginosa*, *P. putida*, and *P. fluorescens*. Residues that may interact with TC are bolded and underlined. The first letter indicates the amino acid found in *P. aeruginosa*, *P. putida*, and *P. fluorescens*. The number indicates the position. The second letter indicates the amino acid in *P. protegens* at the same position.

| **Mutation** | **Location** |
| --- | --- |
| G731A | linking beta-sheets (structural) |
| **A620P** | Loop 7 |
| **S612A** | Loop 7 |
| A483D | Linking beta-sheets (structural) |
| V423T | linking beta-sheets (structural) |
| **D380N** | In proximity to the native ligand binding site |
| H367N | Beta-sheet (structural) |
| A346S | Beta-sheet (structural) |
| Y343F | Beta-sheet (structural) |
| R319H | linking beta-sheets (structural) |
| Q303K | linking beta-sheets (structural) |
| A198T | Alpha-helix within plug domain (structural) |
| S192A | Linking alpha helices within plug domain (structural) |
| A62P | Signalling domain |
| P61A | Signalling domain |
| I60V | Signalling domain |
| P42A | Signalling domain |
| V6I | Signalling domain |

**SUPPLEMENTARY FIGURES**

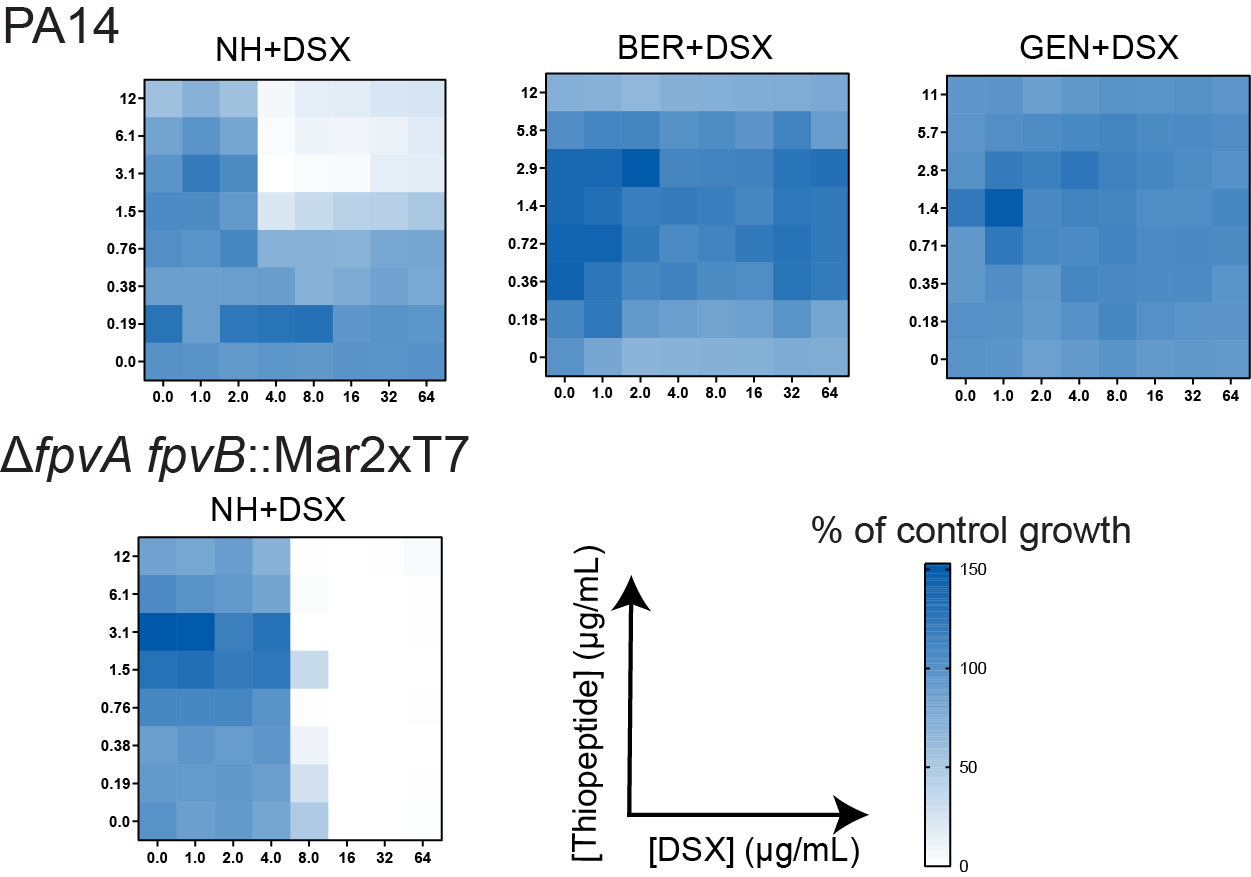

**Supplementary Figure 1.** Some thiopeptides did not synergize with DSX. Assays were conducted in 10:90. Averaged results from three independent experiments are shown. Top row are checkerboards conducted against PA14 with each thiopeptide whereas bottom row are checkerboards conducted against the pyoverdine receptor-null mutant. The concentration of the thiopeptides are indicated along the column (maximum concentration of 10 µM in µg/mL) and the concentration of DSX is indicated along the row (maximum concentration of 64 µg/mL). Growth is proportional to the intensity of blue whereas white indicates no growth. BER and GEN were not tested against the pyoverdine receptor-null mutant due to lack of activity against wild type PA14.

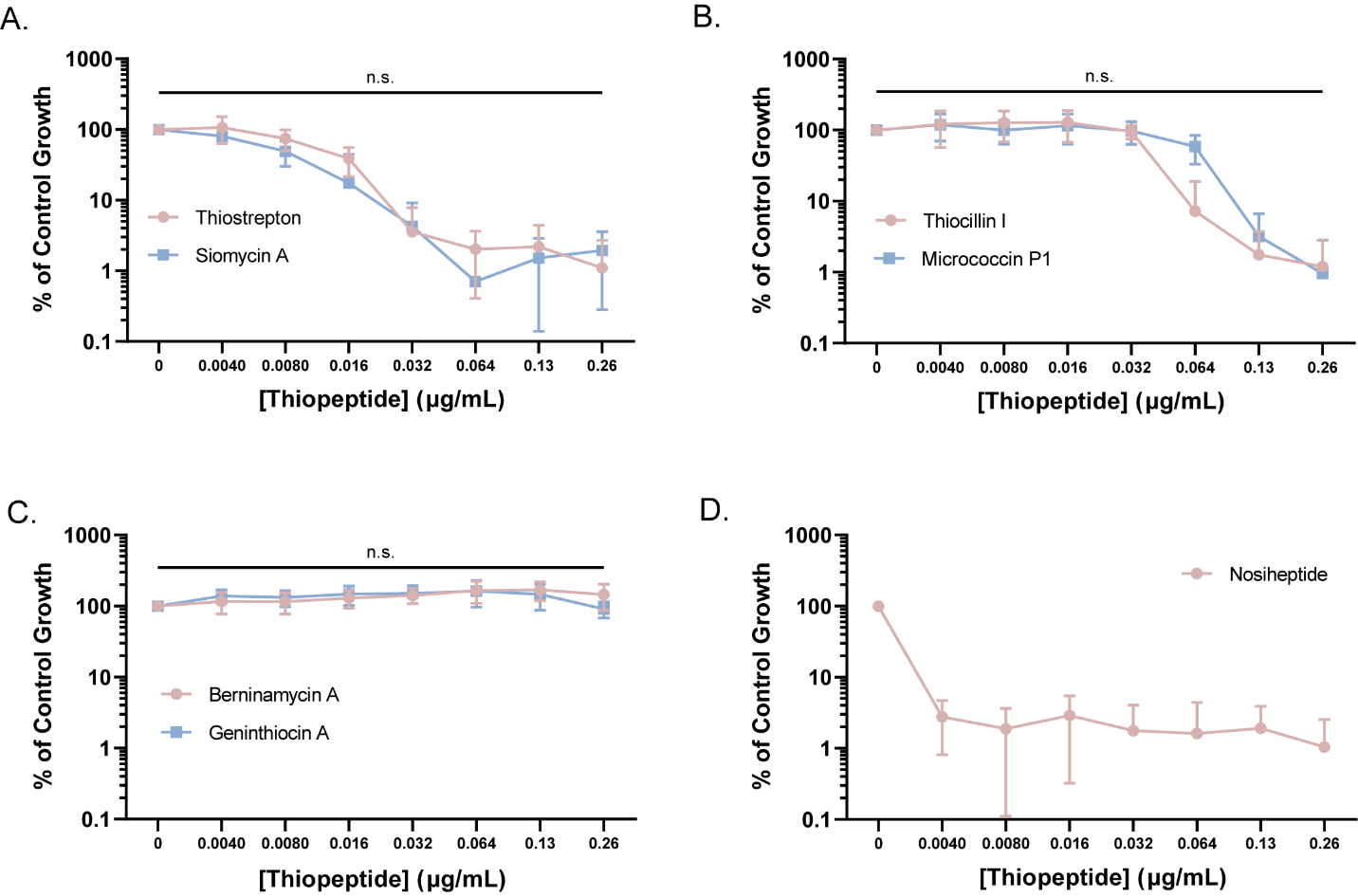

**Supplementary Figure 2.** NH is highly potent against MRSA USA300. Dose-response assays of thiopeptides against MRSA USA300 in 10:90; A) TS and SM, B) TC and MC, C) BER and GEN, D) NH. Assays were done in triplicate and the averaged values are shown. n.s, not significant. Significance was determined using two-way ANOVA followed by Sidak’s multiple-comparison test.

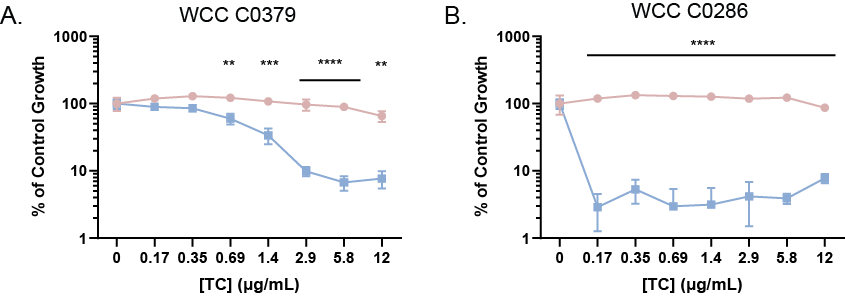

**Supplementary Figure 3.** TC + DSX inhibits growth of TS-resistant *P. aeruginosa* and *A. baumannii* clinical isolates. A) *P. aeruginosa* WCC C0379 and B) *A. baumannii* C0286. Results are averaged from three independent experiments. **, p<0.01; ***, p<0.0005, ****, p<0.0001. Significance was determined using two-way ANOVA followed by Sidak’s multiple-comparison test.

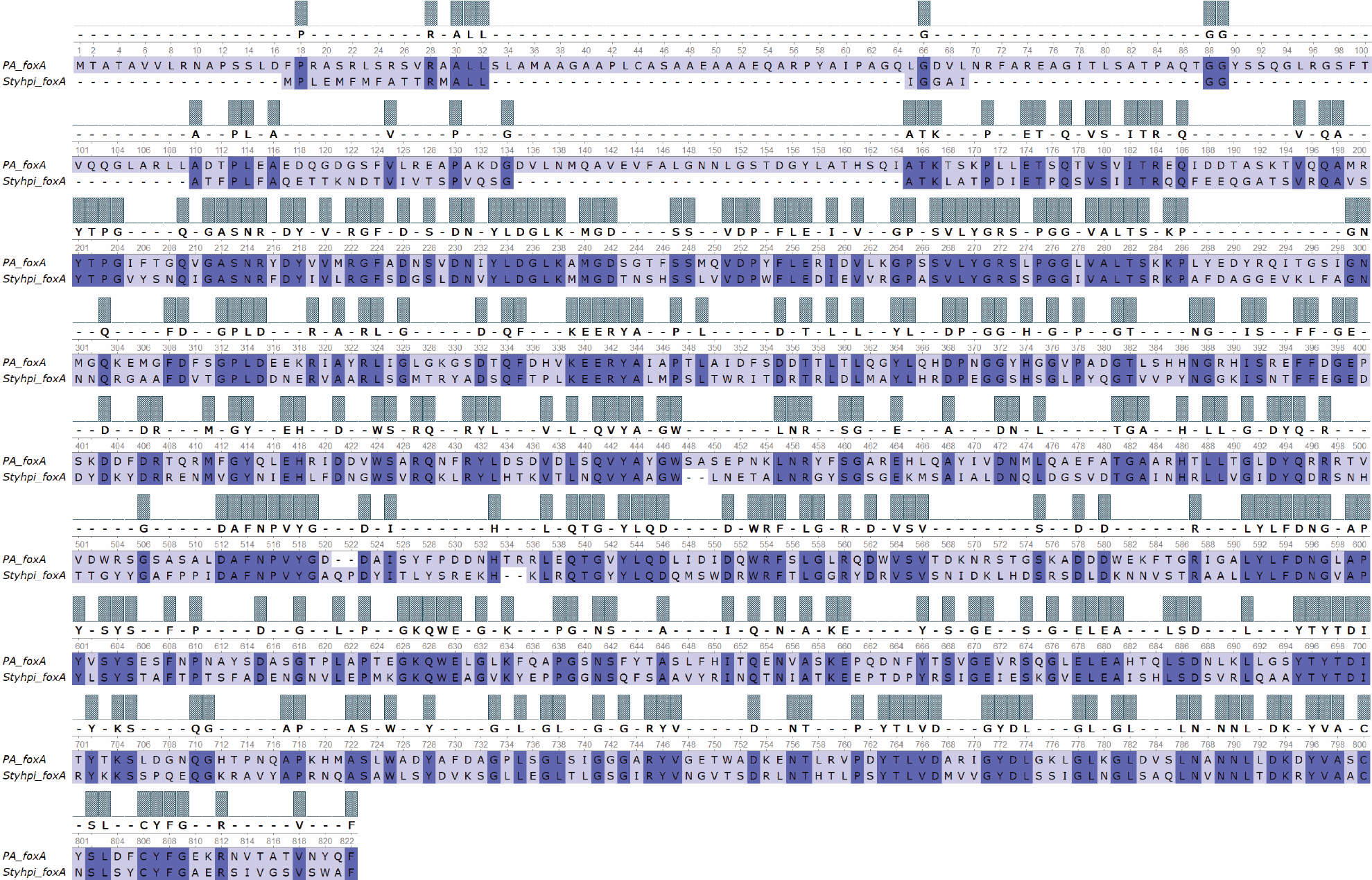

**Supplementary Figure 4.** MUSCLE Alignment of *P. aeruginosa* PAO1 and *S.* Typhimurium SL1344 FoxA. The two proteins share 40% identity. Alignment was done using Unipro UGENE. Accession number for *P. aeruginosa* PAO1 FoxA NP_251156.1. *S.* Typhimurium SL1344 FoxA sequence was obtained from GenBank (Accession number CBW16459.1). Residues involved in ferrioxamine uptake are conserved: Y218, H374, and Q441.

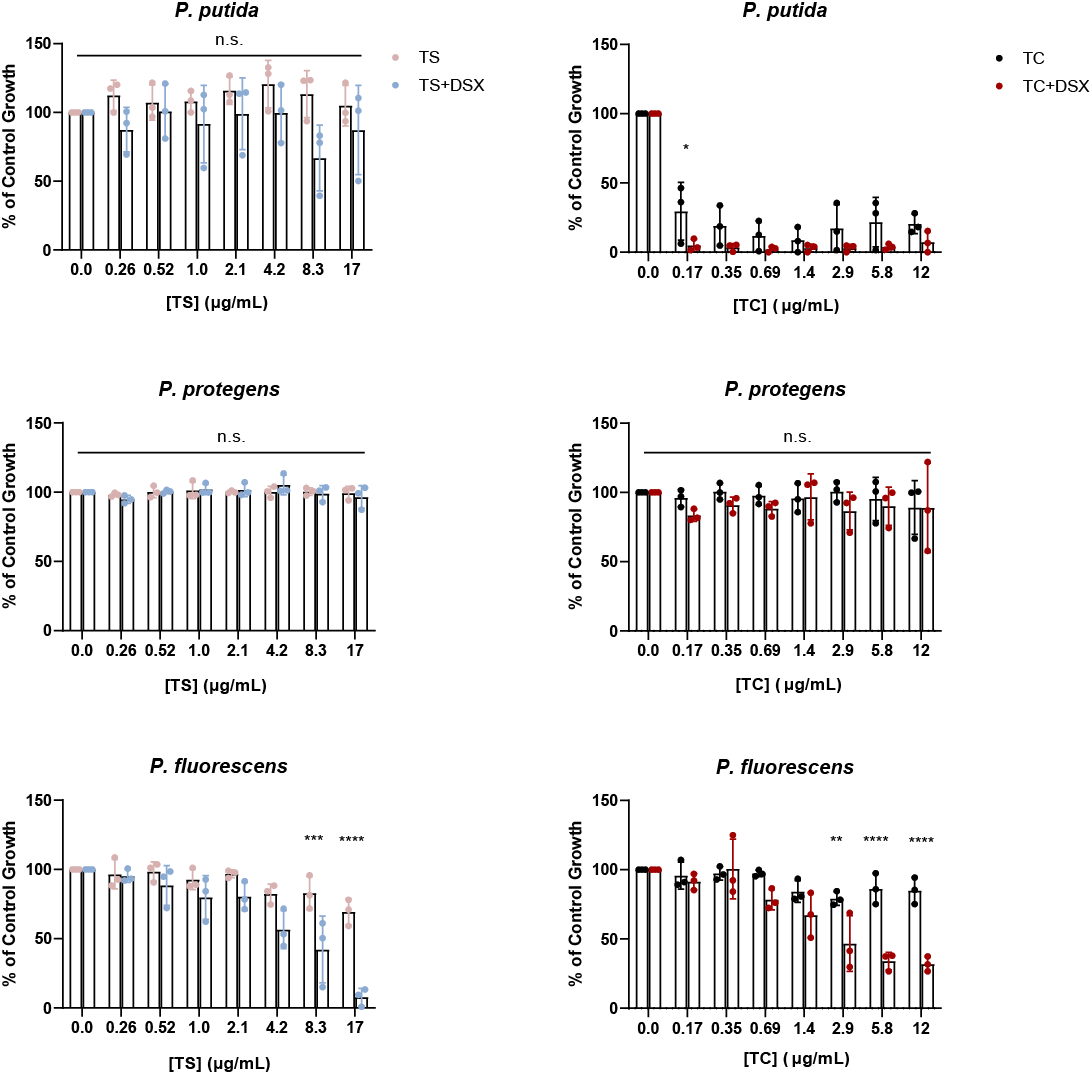

**Supplementary Figure 5.** Susceptibility to TS and TC vary between different *Pseudomonas* species. Dose response assays with TS, TS + DSX, TC, and TC + DSX were tested against *P. putida, P. protegens,* and *P. fluorescens*. The colour of the circles indicates the % of control growth at different concentrations of thiopeptide or thiopeptide plus DSX – TS (pink circles), TS + DSX (blue circles), TC (red circles), and TC + DSX (black circles). n.s., not significant; *, p<0.05; **, p<0.01, ***; p<0.0005; ****, p<0.0001. Results are averaged from three independent experiments and two-way ANOVA was used to determine significance followed by Sidak’s multiple-comparison test.

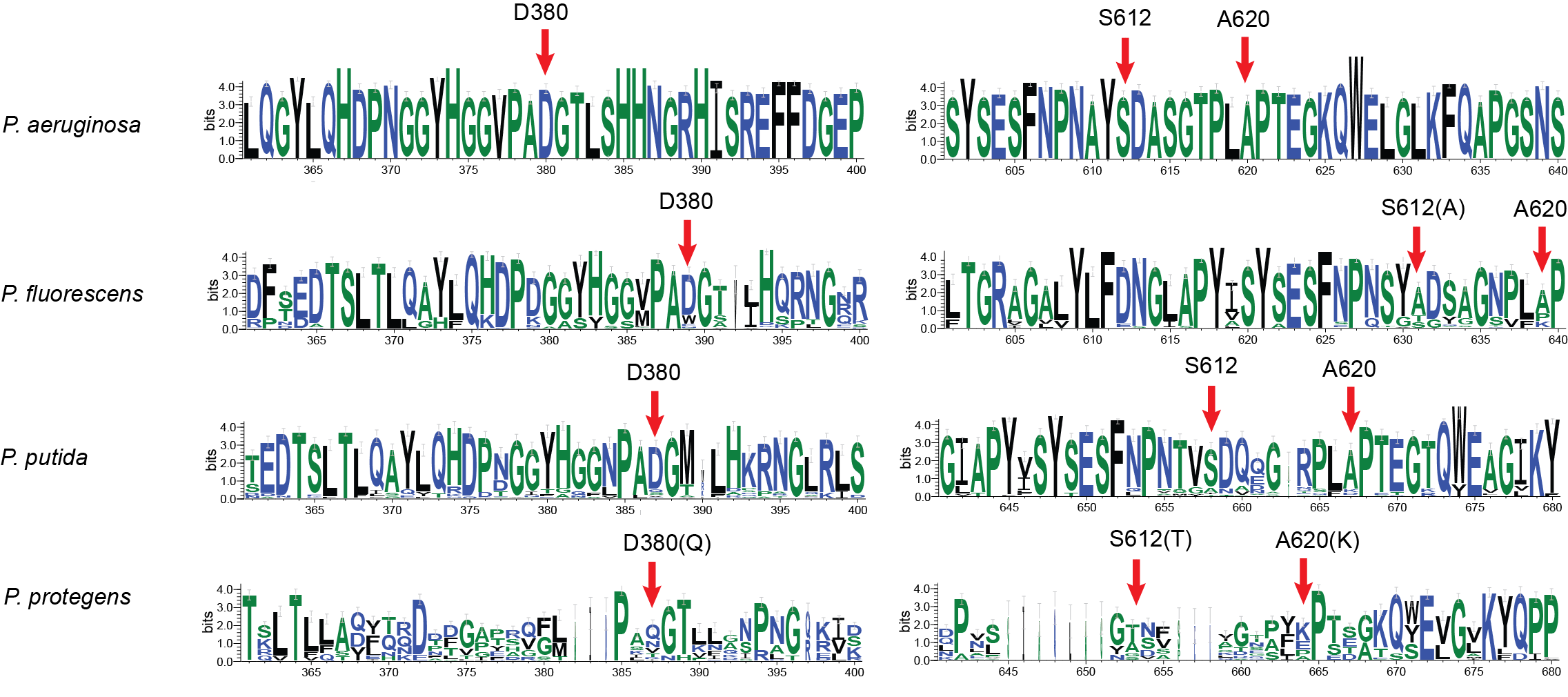

**Supplementary Figure 6.** Species susceptible to TC have a specific pattern of conserved amino acid residues in FoxA. A BLAST search was conducted using *P. aeruginosa* PAO1 FoxA against *P.aeruginosa, P. fluorescens, P. putida* (all TC susceptible)*,* and *P. protegens* (TC resistant)*.* The top 100 hits were aligned (MUSCLE). The consensus sequence was determined for each species and depicted as a sequence logo. Residues considered as potentially important for TC interaction with FoxA were compared to the consensus sequence (Supplementary Table 4). The positions of the residues of interest (i.e. D380, S612, A620) are numbered according to *P. aeruginosa* FoxA. Amino acids in blue are hydrophilic, green are neutral, and black are hydrophobic. Sequence logos were generated using WebLogo.
